# Distinct roles of the human cuneiform and pedunculopontine nuclei in gait initiation and freezing of gait

**DOI:** 10.64898/2026.01.13.699281

**Authors:** Mathieu Yeche, Antoine Collomb-Clerc, Katia Lehongre, Julie Bourilhon, Yannick Mullié, Nathalie George, David Grabli, Huiling Tan, Marco Romanato, Claire Olivier, Hayat Belaid, Brian Lau, Carine Karachi, Marie-Laure Welter

## Abstract

Freezing of gait in Parkinson’s disease (PD) is a major cause of disability, often resistant to dopaminergic therapy and deep brain stimulation (DBS). Its underlying mechanisms remain unclear, mainly because the roles of the human mesencephalic locomotor region (MLR) nuclei are not well understood. Here, we combined rare local field potentials (LFP) recordings from the cuneiform (CuN) and pedunculopontine nuclei (PPN) with biomechanical markers of gait initiation (GI) in four PD patients. We identified functional differences: increases in CuN alpha-band activity precede anticipatory postural adjustments (APA) and correlate with the rhythm of upcoming steps, whereas decreases in PPN beta-band activity occur during APA just before the lead foot lifts off. Imminent freezing is characterized by a breakdown of this organization, marked by mistimed alpha-band surges across the MLR and abnormal PPN beta-band modulation. CuN stimulation selectively improved the stepping rhythm, while PPN stimulation worsened pace or forward vigor. Furthermore, exaggerated mesencephalic alpha-band power was associated with poor clinical responses. These results clarify the individual roles of MLR nuclei in human locomotion and identify pathological alpha dynamics as a biomarker for advancing adaptive neurostimulation.

## INTRODUCTION

Freezing of gait (FOG) is a highly debilitating and poorly understood symptom of Parkinson’s Disease (PD) ^1,2^. It is characterized by a sudden, transient inability to step effectively, leading to falls and loss of independence. It often shows limited response to dopaminergic therapy and routine deep brain stimulation (DBS). Recent anatomical, imaging, and physiological studies emphasize the brainstem mesencephalic locomotor region (MLR) as a critical hub for gait control ^3–6^ and a potential target for DBS in treating gait disorders and falls in PD ^7–10^. However, clinical trials targeting the MLR have yielded inconsistent results ^10–14^, highlighting the urgent need to further understand the functional architecture of this complex region in humans ^15,16^.

The MLR consists of two nuclei, without sharply defined anatomical boundaries: the ventral pedunculopontine nucleus (PPN) and the dorsal cuneiform nucleus (CuN). These nuclei exhibit distinct circuit integration profiles, suggesting divergent functions. The PPN is densely embedded within basal ganglia loops, receiving massive projections from basal ganglia outputs (the substantia nigra pars reticulata and the internal globus pallidus) and primarily integrates sensorimotor and associative information ^17–21^. Its downstream influence is exerted through projections to the pontine reticular formation and ascending feedback loops to the thalamus. In contrast, the CuN receives weaker basal ganglia inputs, which are more restricted to the limbic circuits (e.g., ventral striatum/pallidum). Furthermore, the CuN receives predominant inputs from threat-detection centers ^22,23^, such as the inferior colliculus and the periaqueducal gray (PAG) and sends excitatory projections to the medullary reticular formation (MRF) ^5,17,24^, positioning it to bypass finer postural control circuits and drive rapid, high-speed locomotion ^25^. In mice, although the CuN contains both glutamatergic and GABAergic neurons ^19,21^, optogenetic activation of the glutamatergic population is sufficient to drive high-speed locomotion ^4,26–28^. The PPN comprises an even more heterogeneous population – including cholinergic, glutamatergic, and GABAergic neurons ^6,25,29^– that modulates posture and exploration and can even halt locomotion, depending on the cell type activated ^19,21,29–31^. In non-human primates, tonic and phasic-rhythmic neuronal activity are directly linked to locomotion, and increased burst activity is observed within the caudal MLR following dopaminergic depletion ^32^. Furthermore, lesions of PPN cholinergic neurons cause specific locomotor and postural problems ^33^.

In humans, there is limited direct evidence supporting a specific spatial organization of function ^34^. Intraoperative recordings show that CuN neurons firing during simulated stepping ^35^, and stimulation of this region can induce involuntary rhythmic leg muscle activity ^36^. At the same time, PPN local field potential (LFP) recordings reveal neural modulations during limb movements ^37–39^, imagined gait ^40,41^ and indicate that theta- and alpha-band activity are related to gait control ^42–44^. Additionally, there is increased PPN-cortical alpha coherence during locomotion ^45^. Critically, however, no previous study has recorded from both the CuN and PPN simultaneously to clarify their specific roles. As a result, it remains unknown whether MLR nuclei exhibit distinct electrophysiological signatures during locomotion and how these signatures relate to FOG signature.

This knowledge gap has clinical implications. Current MLR-DBS trials primarily target the PPN, but variability in the target and the unresolved contributions of the CuN versus PPN may explain the limited efficacy in treating FOG ^9,11–13,46,47^. Furthermore, because FOG is an episodic phenomenon, standard continuous stimulation may fail to address the dynamic neural dysfunction preceding freezing episodes, which can be detected during gait initiation (GI) ^48^. To develop effective therapies, we must identify the specific temporal dynamics of MLR nuclei ^49^ during the transition from standing to walking, specifically distinguishing between anticipatory postural adjustments (APAs), which prepare balance, and the rhythmic stepping phase.

In this study, we used DBS electrodes in the MLR, with contacts in both the CuN and PPN, to record LFPs in four PD patients with dopa-resistant FOG. By combining these recordings with biomechanical analyses of GI, we tested the hypothesis that the CuN and PPN have distinct, time-specific roles in human gait control. We identified electrophysiological signatures of impeding freezing, encoded in specific frequency bands around GI, which predicted gait quality under DBS. These findings provide a mechanistic understanding of FOG development and lay the foundation for next generation adaptative DBS strategies.

## RESULTS

### Specific CuN and PPN neuronal activity

We recorded LFPs from the MLR of four patients with advanced PD during self-initiated gait, alongside force platform, motion capture, and electromyography (**Fig. 1a**, **Supplementary Table 1, Supplementary Fig. 1a-b**). A total of 56 bipolar LFP recordings (7 per electrode; one electrode per hemisphere) were localized using a histology-based human brainstem atlas ^13,50^. Of these, 13 dipoles were localized in the PPN (cholinergic population) and 15 in the CuN, with each patient contributing at least one dipole in both nuclei (**Fig. 1b**).

**Fig. 1.**
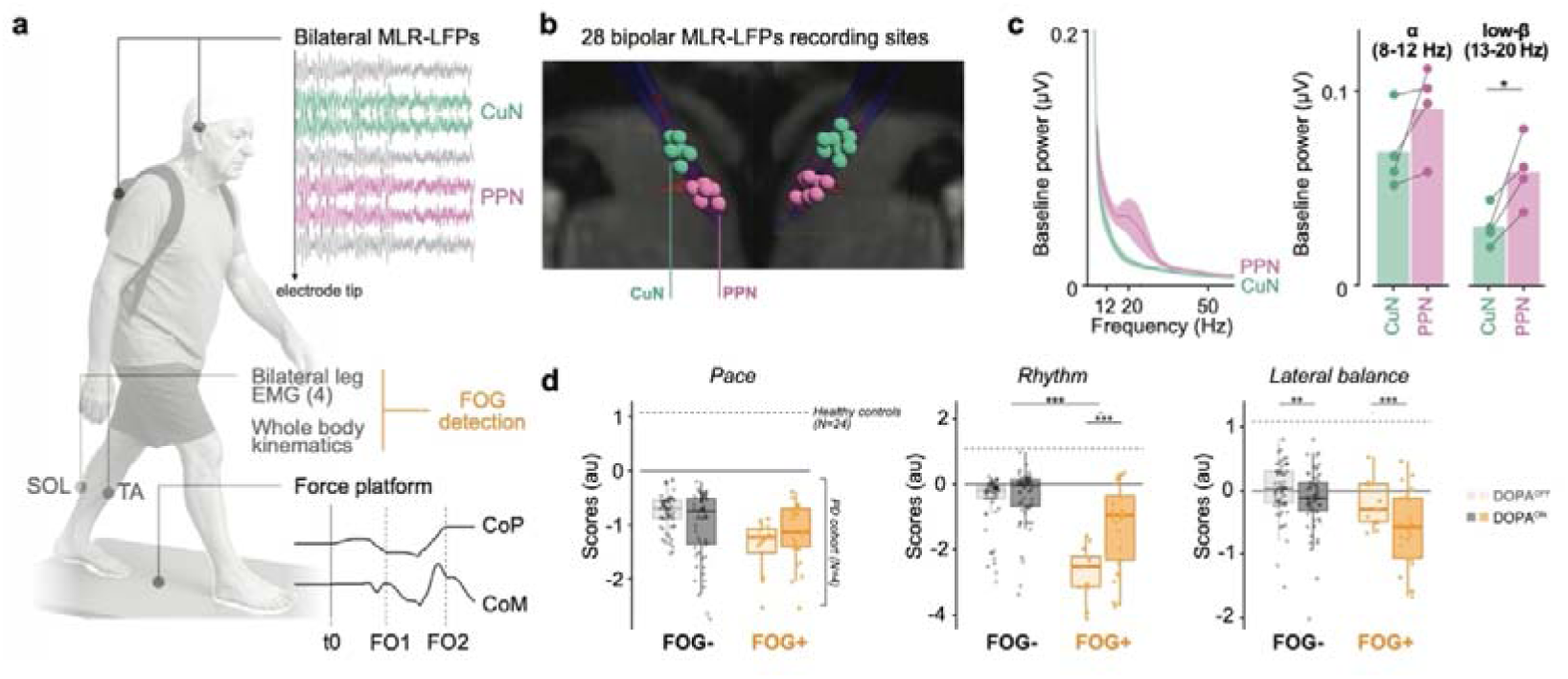
Multimodal profiling of MLR dynamics during gait initiation. **a.** Schematic of the experimental setup combining simultaneous MLR-LFP recordings (via externalized DBS leads) with whole-body motion capture, force platform dynamometry (Center of Pressure, CoP; Center of Mass, CoM), and surface electromyography (EMG) of soleus (SOL) and tibialis (TA). Temporal events are defined as: t0 (onset of anticipatory postural adjustment phase, APA), FO1 (foot-off, first step) and FO2 (foot-off, second step). **b.** Anatomical reconstruction of recording sites. Sagittal view of DBS electrodes superimposed on a stereotaxic brainstem atlas. The red area demarcates the putative cholinergic PPN. LFP bipolar derivations are color-coded by location: cholinergic PPN (n=13, purples) and dorsal non-cholinergic CuN (n=14, green). **c.** Baseline mesencephalic power spectra. Left: Mean power spectral density (PSD) at rest for CuN and PPN. Right: Comparison of average baseline power in the alpha (8–12 Hz) and low-beta (13–20 Hz) bands. Dots indicate patient-level averages across contacts (n = 4 patients). *p□<□0.05, paired t-test. **d.** Gait initiation mechanics with and without freezing. Box plots showing kinetic sub-scores, i.e. *Pace*, *Rhythm*, and *Lateral balance*, for trials with (FOG+) and without (FOG−) freezing episodes, in DOPA^OFF^ and DOPA^ON^ states. ***p<0.001, LMM, Bonferroni corrected. Box elements: center line, median; box limits, upper and lower quartiles; whiskers, 1.5 times interquartile range; dots, individual trials.

Baseline (standing still) activity showed higher low-beta (13–20 Hz) power in the PPN than in CuN (t_(3)_ = –3.46, *p* = 0.041; **Fig. 1c**), whereas alpha (8–12 Hz) power did not differ significantly (t_(3)_ = –2.00, *p* = 0.14; **Fig. 1c**). Under dopaminergic medication, baseline MLR LFP activity was not significantly modified (e.g., beta power: t_(3)_ = 0.86, *p* = 0.45), although one participant showed a marked reduction in PPN beta-band activity DOPA^ON^ vs DOPA^OFF^ conditions (**Supplementary Fig. 2a**). During GI, dopaminergic medication did not induce significant changes in MLR alpha or beta power (**Supplementary Fig. 2b,c**), with a smaller theta increase during APAs in the CuN (DOPA^OFF^ vs. DOPA^ON^: 0.13 to 0.64 s, β = 1.85 ± 0.48 dB, *p* = 0.0001, **Supplementary Fig. 2b,c**). Given the absence of alpha and beta modulations between dopaminergic medication conditions, we pooled LFP data from both DOPA states in subsequent analyses, keeping medication condition as a covariate.

### Gait initiation is altered by FOG imminence

All four PD patients experienced FOG during the session, yielding 38 FOG trials (FOG+; 12 DOPA^OFF^, 26 DOPA^ON^), with FOG occurring on average 13.5 steps after GI (IQR, 10–23), during walking or turning. We also collected 115 trials without FOG episodes (FOG−; 57 DOPA^OFF^, 58 DOPA^ON^).

We compared GI parameters across FOG+ and FOG- trials, and between DOPA^OFF^ and DOPA^ON^ conditions using principal component scores: *Pace* (correlating with step length, propulsion and stepping vigor), *Rhythm* (correlating with step timing and cadence), and *Lateral balance* (correlating with step width and lateral center of mass velocity) (**Supplementary Fig. 1c**). *Pace* was unaffected by either FOG imminence (β = 0.0039 ± 0.068 a.u., *p_c_*= 0.99) or dopaminergic medication (β = 0.047 ± 0.056 a.u., *p_c_* = 0.99, **Fig. 1d**). *Rhythm*, by contrast, was significantly impaired when FOG was imminent (FOG+ vs FOG-trials; β = 0.63 ± 0.16 a.u., *p_c_*< 0.001), with a stronger effect in the DOPA^OFF^ state (DOPA^OFF^ vs DOPA^ON^; β = 1.08 ± 0.28 a.u., *p_c_*< 0.001; **Fig. 1d**). *Lateral balance* did not significantly differ between FOG+ and FOG- trials (FOG+ vs FOG- trials; β = -0.057 ± 0.086 a.u., *p_c_* > 0.99) but was significantly deteriorated under dopaminergic medication (β = 0.40 ± 0.07 a.u., *p_c_*< 0.001), with a more pronounced effect in FOG+ trials (β = –0.33 ± 0.15 a.u., *p_c_* = 0.026; **Fig. 1d**). In FOG+ trials, dopaminergic medication exerted opposite effects, enhancing *Rhythm* (-1.07 ± 0.24 a.u., *p_c_* < 0.001) while worsening *Lateral balance* (0.56 ± 0.13 a.u., *p_c_* < 0.001; **Fig. 1d**).

Together, these findings demonstrate that impaired *Rhythm* and *Lateral balance* during GI are key markers of FOG imminence in PD. Dopaminergic medication amplified their dissociation by rescuing *Rhythm* but impairing *Lateral balance*.

### Specific contribution of CuN and PPN neural dynamics during GI

Examining LFP activity during GI, irrespective of FOG imminence, CuN and PPN exhibited distinct patterns (**Fig. 2a**). In the CuN, alpha-band power increased before APAs onset. In the PPN, pronounced beta-band desynchronization occurred during APAs, with alpha-band activity rising afterward (during gait execution) (**Fig. 2a**). Direct time-frequency comparisons confirmed these differences: the pre-APA alpha increase was stronger in the CuN (-0.35 s to 0.04 s, β = 1.22 ± 0.36 dB, *p* = 0.0026), whereas post-APA low-beta desynchronization was stronger in the PPN (-0.17 s to 0.40 s, β = 0.88 ± 0.25 dB, *p* = 0.0012; **Fig. 2b,c**).

**Fig. 2.**
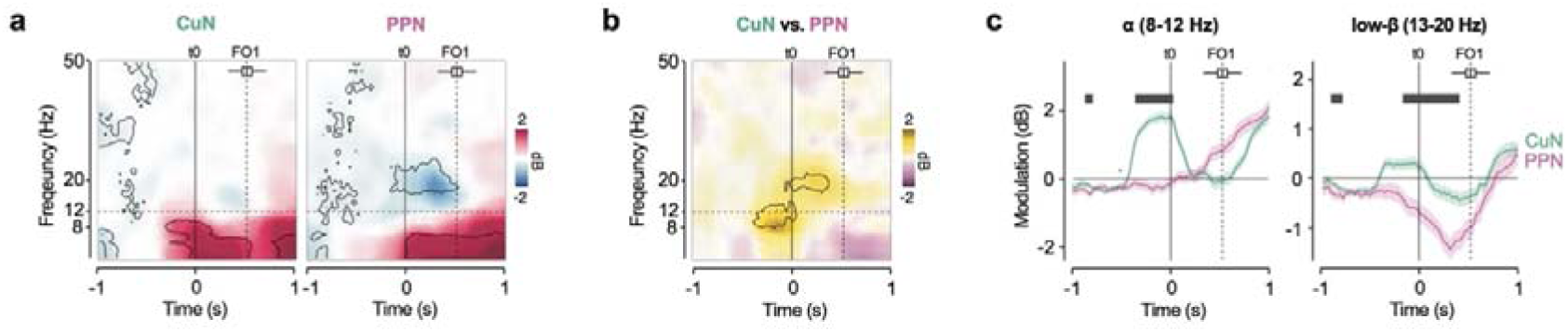
Spatiotemporal dissociation of CuN and PPN activity during gait initiation. **a.** Time–frequency analysis of CuN and PPN activity. Spectrograms show event-related power modulations (relative to baseline) aligned with the onset of anticipatory postural adjustments (t0). The dashed vertical line indicates the average time of the first foot-off (FO1); the horizontal boxplot above illustrates the distribution of FO1 timings across all trials. Black contours mark significant clusters (p□<□0.05, LMM). **b.** Functional contrast map. Time-frequency plot illustrating the direct statistical comparison between PPN and CuN power modulation (PPN – CuN). Warm colors indicate higher power in CuN; cool colors indicate higher power in PPN. Significant differences are outlined in black (p < 0.05, LMM). **c.** Frequency-specific temporal dynamics. Averaged power modulation time-courses for the alpha (8–12 Hz) and low-beta (13–20 Hz) bands in the CuN (green) and PPN (purple). Black bars at the top indicate time points with significant divergence between the two nuclei (p < 0.05, LMM). Shaded areas represent standard error of the mean (SEM).

### FOG imminence dissociates CuN and PPN neural encoding of GI

To further understand how MLR neuronal activity is shaped by FOG imminence, we compared LFP between FOG+ and FOG− trials, in both nuclei (**Fig. 3**). When FOG was imminent, pre-APA alpha-band power tended to decrease in the CuN (FOG+ vs. FOG- trials, -0.3 s to 0 s, β = 0.39 ± 0.39, p = 0.3132, **Fig 3a-c**). Conversely, per-APA alpha-band power significantly increased in both the CuN and PPN for FOG+ trials (FOG+ vs. FOG- trials, 0.04 s to 0.67 s, β = 1.09 ± 0.29 dB, *p* = 0.0002, **Fig. 3a,b**) with no significant differences between MLR subregions (FOG+ trials: CuN vs PPN, 0.04 s to 0.67 s, β = 0.11 ± 0.56 dB, *p* = 0.84, **Fig 3b,c, Supplementary Fig. 3**). In the PPN, low-beta desynchronization was significantly stronger for FOG+ trials compared to FOG- trials (0.43 s to 0.67 s, β = 1.09 ± 0.44 dB, *p* = 0.0137; **Fig. 3a-c**).

**Fig 3.**
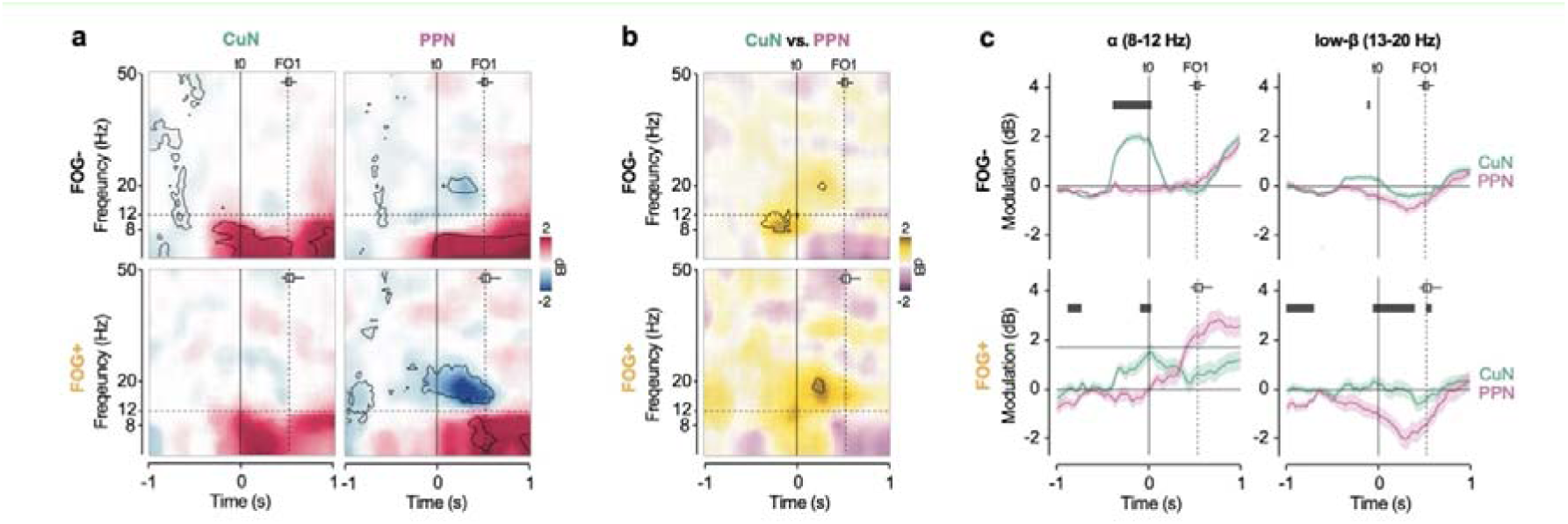
Mesencephalic neural signatures of imminent freezing. **a.** State-dependent spectral dynamics. Time-frequency maps showing LFP power modulation (relative to baseline) in the CuN and PPN, separately for effective gait initiation without subsequent freezing (FOG−, upper row) and trials where successful gait initiation led to freezing (FOG+, lower row). Data are aligned to the APA onset (t0). Black contours indicate significant event-related modulation (p < 0.05, LMM). **b.** Collapse of functional segregation. Time-frequency plots showing the difference in power modulation between PPN and CuN (PPN - CuN) during FOG− and FOG+ trials. Significant spatial dissociation (black contours, p < 0.05) is prominent in FOG− trials but diminishes in FOG+ trials. **c.** Frequency-specific breakdown. Temporal profiles of alpha (8-12 Hz) and low-beta (13-20 Hz) power modulation in CuN (green) and PPN (purple) for FOG− versus FOG+ conditions. Black bars indicate significant differences between nuclei (p < 0.05, LMM). Shaded areas represent SEM.

Looking at the correlation between MLR-LFP and GI performance, we found that CuN alpha-band power at APA onset significantly positively correlated with *Rhythm* and *Lateral balance* scores (*Rhythm* : -0.23 s to 0.16 s, β = 0.73 ± 0.21 dB, *p* = 0.0005; *Balance*: -0.08 s to 0.19 s, β = 1.33 ± 0.39 dB, *p* = 0.0006, **Fig. 4)**. In the PPN, alpha-band power at APA onset negatively correlated with *Pace* (-0.11 s to 0.16 s, β = -1.50 ± 0.47 dB, *p* = 0.002), and with *Rhythm* at the end of APA (time of FO1, 0.43 s to 0.70 s, β = –0.90 ± 0.30 dB, *p* = 0.0041; **Fig. 4b**).

**Fig. 4.**
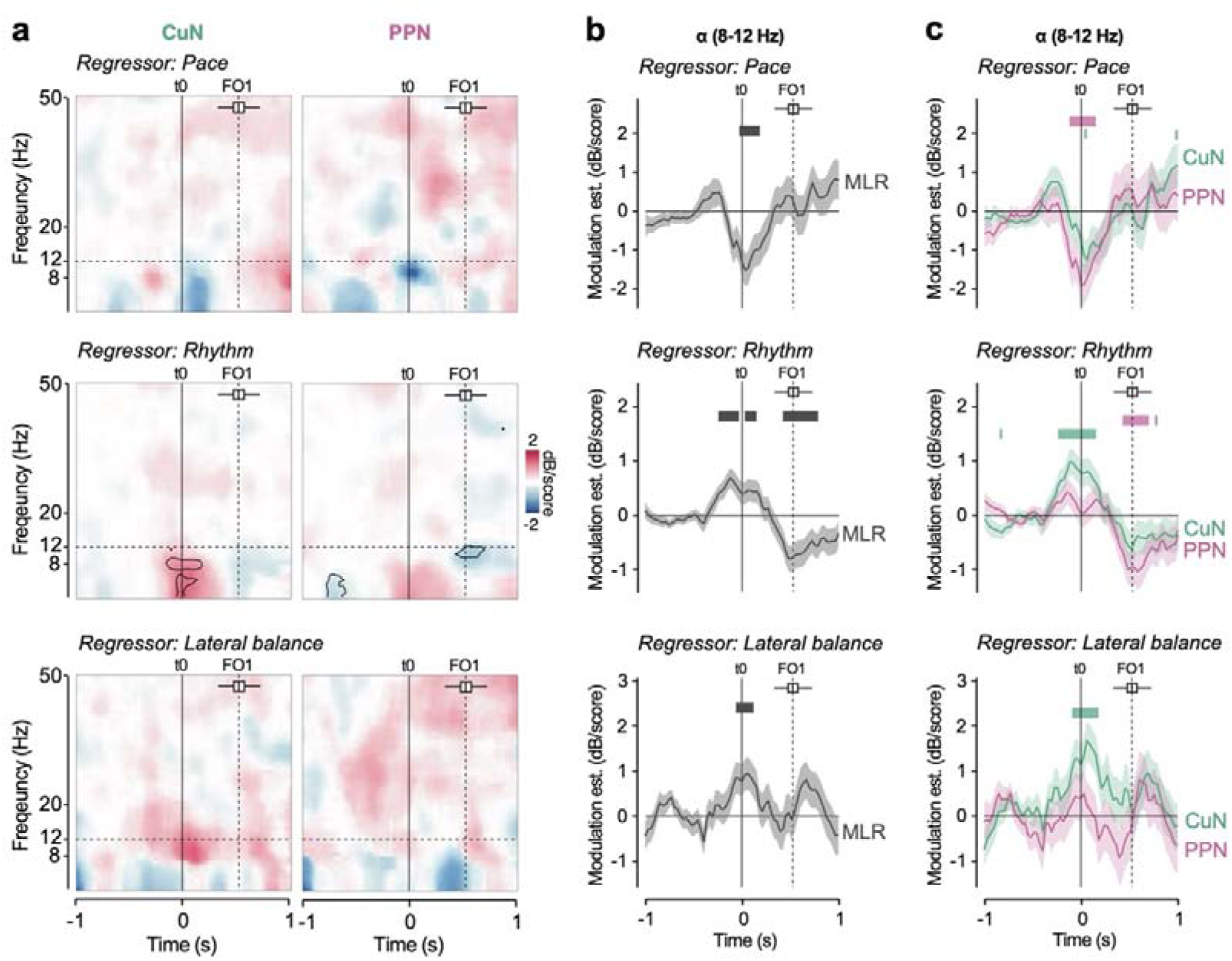
Alpha-band dynamics encode gait initiation quality. **a.** Spectral correlates of gait mechanics. Time-frequency maps display the statistical relationship (LMM coefficients) between LFP power in CuN (left column) and PPN (right column) and biomechanical scores: *Pace*, *Rhythm*, and *Lateral Balance*. Data are aligned to APA onset (t0). Warm colors indicate positive correlations; cool colors indicate negative correlations. Black contours outline significant clusters (p < 0.05, LMM). The horizontal boxplot above shows the distribution of first foot-off (FO1) times; the dashed vertical line marks the mean FO1. **b.** Alpha-band specific encoding across MLR. Temporal profiles of the correlation between alpha power (8–12 Hz) and biomechanical scores. Black bars denote time points where the correlation strength significantly differs from zero (p < 0.05, LMM). Shaded areas represent SEM. **c.** Alpha-band specific encoding for CuN and PPN. Temporal profiles of the correlation between alpha power (8–12 Hz) and biomechanical scores for CuN (green) and PPN (purple). Green and purple bars indicate time points during which the correlation is significant for CuN and PPN, respectively (p < 0.05). Black bars denote time windows where the correlation strength differs significantly between nuclei (p < 0.05, LMM). Shaded areas represent SEM.

Together, these results indicate that early CuN alpha activity supports *Rhythm* and *Balance* at GI, but exaggerated alpha activity during APAs phase across the MLR disrupts GI with FOG imminence, with additional lowered low-beta PPN activity.

### CuN-DBS improves Rhythm in relationship with alpha band power

We assessed the effects of DBS of the CuN (DBS^CuN^) or PPN (DBS^PPN^) relative to no stimulation (DBS^OFF^) using a double-blind, randomized design (**Fig. 5a,b**).

**Fig. 5.**
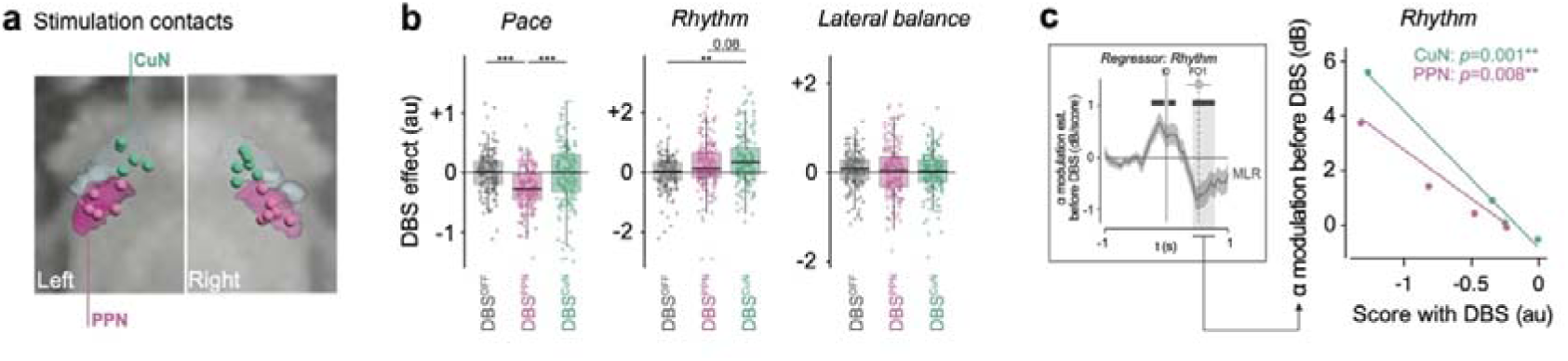
Mesencephalic alpha activity predicts differential response to MLR-DBS. **a.** Anatomical localization of active contacts. Reconstruction of active DBS contacts used for CuN (green) and PPN (purple) stimulation, superimposed on a stereotaxic atlas. The purple area demarcates the putative cholinergic PPN; the light green area corresponds to the dorsal non-cholinergic CuN. **b.** Differential clinical effects. Box plots showing biomechanical gait initiation scores (*Pace*, *Rhythm*, and *Lateral Balance*) under three conditions: DBS^OFF^, DBS^CuN^ and DBS^PPN^ conditions (data pooled across DOPA^OFF^ and DOPA^ON^ states). CuN stimulation selectively improves *Rhythm*, whereas PPN stimulation worsens *Pace*. Box elements: center line, median; box limits, upper and lower quartiles; whiskers, 1.5 times interquartile range; dots, individual trials. **p < 0.01, ***p < 0.001, LMM, Bonferroni-corrected. **c.** Alpha-band activity as a predictor of clinical outcome. Regression plots showing the relationship between the *Rhythm* score under DBS and the amplitude of alpha-band power during gait initiation (measured without DBS). Stronger alpha modulation predicts worse *Rhythm* outcomes, suggesting it marks resistance to therapy.

DBS^PPN^ significantly worsened *Pace* compared with both DBS^OFF^ and DBS^CuN^ (DBS^PPN^ vs. DBS^OFF^: β = 0.30 ± 0.04, *p_c_* < 0.001; DBS^PPN^ vs. DBS^CuN^: β = –0.20 ± 0.04, *p_c_* < 0.001; **Fig. 5b**), whereas DBS^CuN^ had no significant effect (DBS^CuN^ vs. DBS^OFF^, β = 0.10 ± 0.04, *p_c_* = 0.11). In contrast, DBS^CuN^ significantly improved *Rhythm* compared with DBS^OFF^ (DBS^CuN^ vs. DBS^OFF^: β = –0.27 ± 0.08, *p_c_* = 0.0023; **Fig. 5b**), whereas DBS^PPN^ had no significant effect (DBS^PPN^ vs. DBS^OFF^ β = –0.086 ± 0.076, *p_c_* = 1.00; DBS^PPN^ vs. DBS^CuN^: β = –0.18 ± 0.08, *p_c_* = 0.083; **Fig. 5c**). *Lateral balance* was not significantly affected by either DBS^PPN^ or DBS^CuN^ (DBS^PPN^ vs. DBS^OFF^: β = 0.073 ± 0.066, p = 1.00; DBS^PPN^ vs. DBS^CuN^: β = 0.042 ± 0.066, p = 1.00; **Fig. 5b**).

Across patients, alpha increases during APAs were negatively correlated with *Rhythm* under both DBS^CuN^ (β = –4.26 ± 0.59 dB/au, *p* = 0.0001) and DBS^PPN^ (β = –3.14 ± 0.69 dB/au, *p* = 0.0019) conditions (**Fig. 5c)**. This relationship was significant in both FOG+ (β = –5.21 ± 0.64 dB/au, *p* < 0.0001) and FOG− trials (β = –2.18 ± 0.64 dB/au, *p* = 0.0094). These findings indicate that exaggerated alpha activity predicted a limited MLR DBS benefit for *Rhythm*.

## DISCUSSION

Our findings show that the human MLR is functionally divided, with distinct neural dynamics in the CuN and PPN nuclei linked to specific components of gait initiation. By combining multisite LFP recordings with kinetic analysis and stimulation, we found that CuN alpha-band activity is timed to trigger APAs that lead to rhythmic stepping, whereas PPN alpha-band activity is time-locked to the onset of stepping. Imminent FOG was characterized by disruption of this spatial and temporal organization with abnormal alpha synchronization and compensatory yet ineffective beta modulation. Importantly, we provide causal evidence that CuN-DBS selectively improves stepping rhythm. At the same time, PPN-DBS worsens forward vigor, challenging current treatment approaches and highlighting alpha dynamics as a key biomarker for adaptive neurostimulation.

### CuN alpha dynamics as the gait rhythm generator

In our study, CuN activity was distinguished from PPN activity by an early alpha-band peak that began well before APAs, decreased during APAs, and then increased again at liftoff of the lead foot. The pre-APA peak positively correlated with the subsequent stepping rhythm, suggesting that the CuN alpha act as a neural trigger for initiating locomotion. The late surge in alpha-band activity, marking the start of stepping, may relate to online locomotor control, consistent with MLR recordings linking theta-alpha power to walking speed in mice ^51^ and humans ^43,44^. While PPN shows no pre-APA alpha-band peak, it does show a surge in alpha-band activity at the start of stepping. However, this timing pattern breaks down when freezing is imminent (i.e., after successful gait initiation by 13.5 steps): the distinctive CuN alpha activity pattern is weakened and associated with a significant deterioration of the stepping rhythm. At the same time, the rise in PPN alpha-band activity no longer marks the start of stepping, indicating a loss of spatial and temporal specificity within the MLR. The disruption in the timing between CuN and PPN neural activity may reflect a failure to effectively transition from APAs to step initiation. We suggest that this loss of spatial and temporal precision in alpha-band activity could serve as an electrophysiological marker of early behavioral indicators of FOG ^44^, such as irregular cadence and shorter step duration ^48,52,53^. It may also relate to a functional decoupling between the CuN and cortical SMA-premotor networks ^54^, which disrupts the essential alignment of postural preparation ^55–57^. Consequently, impaired weight-shifting and faulty step tuning – some of the earliest signs of impeding FOG episodes ^58,59^ – may occur, especially during cognitively demanding gait tasks that increase cortical engagement ^60,61^.

### PPN beta dynamics

In contrast to the CuN, the PPN showed significant low-beta desynchronization during APAs, mirroring beta suppression reported in the MLR during imagined gait ^41^, voluntary leg movements ^62^, and gait-phase transitions ^45^. Anatomically, the PPN is more densely integrated with basal ganglia circuits than the CuN ^21^, and displays functional beta-band coupling with the subthalamic nucleus (STN) ^63,64^. Similar decreases in beta power are observed within STN–cortical circuits during leg movements ^65^, stepping in place ^66,67^, gait initiation ^48^, and walking ^68^, where they correlate with motor readiness and movement vigor^69^. Our findings thus support a pro-kinetic role of PPN beta suppression in facilitating postural readiness and step execution. Unexpectedly, imminent FOG was associated with exaggerated low-beta desynchronization in the PPN. While high beta-band power is typically associated with antikinetic states in the SMA-premotor cortical areas and the STN ^70–73^, recent research shows that increased low-beta STN modulation exists in FOG-prone patients ^48^, which may reflect increased recruitment of prefrontal-medial cortical circuits to compensate for reduced gait automaticity and impaired postural control ^72,74,75^. We propose that the excessive PPN beta modulation observed here represents a similar compensatory response – an attempt to activate postural mechanisms that promote successful gait initiation but ultimately fails to produce effective stepping due to the concurrent alpha-band blockade in the CuN.

### Dissociating clinical effects: CuN versus PPN stimulation

Our double-blind stimulation results provide causal support for the functional dissociation observed in our recordings. CuN-DBS selectively improved gait rhythm, consistent with animals studies ^5,76^, where optogenetic glutamatergic CuN activation initiates and drives high-speed locomotion with physiological stepping patterns in healthy animals ^4,19,29,77,78^, and restores locomotion in Parkinsonian mice ^30,79^. This effect likely relies on the rhythmic activation of descending reticulospinal projections ^17,24,80,81^. Conversely, PPN-DBS paradoxically worsened pace, reducing step length and velocity. This adverse effect may be related to the function of specific PPN populations which can halt locomotion or restrict it to slow, exploratory modes ^19,25,51,82^.

These divergent outcomes may explain the inconsistency observed in previous MLR-DBS trials, which predominantly targeted the PPN ^9,11,83^. ^46,83^. Our findings suggest that the therapeutic efficacy of MLR-DBS on FOG depends on selectively engaging the CuN’s rhythmogenic drive while avoiding the PPN’s braking signals ^36^. This is supported by recent evidence that unilateral CuN stimulation promotes locomotor recovery in rodent models ^84,85^ and human spinal cord injury ^86^.

### Implications for Adaptive DBS

We also identified a predictor of therapeutic resistance: exaggerated mesencephalic alpha-band synchronization. This pathological signature was present across both nuclei and mirrored abnormalities observed in the STN ^48^, pointing to a network-wide collapse of gait automaticity. This finding opens the door to next-generation adaptive DBS (aDBS). While closed-loop systems targeting beta power in cortical-basal ganglia networks show primes in PD ^87,88,89,90^, our data suggest that preventing the pathological alpha surge in the MLR could be a more specific target for alleviating freezing ^49^. By dynamically modulating stimulation based on real-time alpha dynamics, aDBS could theoretically restore the temporal precision of gait initiation without the side-effect of continuous stimulation.

### Limitations and perspectives

The rarity of human MLR recordings and the small number of patients inevitably limit generalizability. However, the consistency of spectral patterns across trials and their alignment with cortical and animal literature strengthen our conclusions. Future studies should expand these findings in animal models and incorporate multimodal recordings across basal ganglia, brainstem, and cortical networks to better characterize the distributed neural dynamics of locomotion and its dysfunction in PD. Given the episodic, context-dependent nature of FOG and its sensitivity to stress or environmental triggers ^1,29,54,91^, the role of limbic and sensory inputs to the MLR also merits further investigation ^22,23,92^.

## Conclusion

We provide a comprehensive dissection of the human MR, revealing that the CuN and PPN play complementary yet distinct roles in initiating gait. We identify aberrant alpha synchronization as a key neural mechanism underlying freezing and a predictor of DBS resistance. These findings call for a re-evaluation of surgical targets in the brainstem and establish a physiological framework for developing refined, adaptive neurostimulation therapies to restore mobility in PD.

## MATERIAL AND METHODS

### Participants

Six patients with PD were enrolled in a randomized, cross-over controlled trial assessing the efficacy of MLR-DBS for gait and balance disorders, including FOG (see ^13^for details). Eligibility criteria were: (1) diagnosis of idiopathic PD according to the United Kingdom Parkinson’s Disease Society Brain Bank criteria; (2) age 18–70 years; (3) presence of gait and/or balance impairment not responsive to dopaminergic treatment, defined as a score >1 on the Movement Disorder Society–Unified Parkinson’s Disease Rating Scale (MDS-UPDRS) part II (item 2.12, walking/balance, and/or item 2.13, freezing of gait) and/or part III (item 3.11, freezing of gait, and/or item 3.12, postural instability) ^93^ in the ON medication state; (4) levodopa responsiveness >40% for other motor symptoms; (5) no contraindication to MRI or DBS surgery; (6) absence of dementia (Mattis Dementia Rating Scale score >129); (7) stable antiparkinsonian medication for ≥3 months before enrolment; (8) willingness to participate; and (9) written informed consent and social security affiliation. The study was conducted in accordance with the Declaration of Helsinki and Good Clinical Practice guidelines. Approval was obtained from the local ethics committee (Comité de Protection des Personnes, Ile-de-France VI), and the trial was registered at ClinicalTrials.gov (NCT02931097).

### Surgical procedure and anatomical localization of recordings

At baseline, patients underwent a comprehensive clinical evaluation before bilateral implantation of MLR-DBS electrodes. MLR targeting was guided by preoperative 1.5T and 3T T1-weighted MRI and an in-house 3D histological deformable atlas of the human brainstem. Intraoperative microelectrode recordings and X-ray fluoroscopy were performed to refine targeting. Bilateral 8-contact DBS leads (DB-2201, diameter 2 mm, Boston Scientific) were implanted and temporarily connected to externalized cables to allow LFP recordings, which were acquired 1–4 days postoperatively before the pulse generator (Vercise PC, Boston Scientific) was implanted (Figure 1a). A 3D helical CT scan obtained the day after surgery was co-registered with the preoperative T1-weighted MRI and overlaid on the MLR histological atlas to confirm lead position and rule out complications. Recording dipoles were defined as the midpoint between each contact pair. Fifteen dipoles were assigned to the CuN and 13 to the PPN subregions and included in subsequent analyses (**Fig. 1b**).

### Gait initiation recordings

Patients underwent biomechanical and physiological gait assessments across four sessions in the Paris Brain Institute Physiology and Analysis of Movement (PANAM) core facility. In each session, patients were tested in two conditions: without dopaminergic medication (DOPA^OFF^, following a 12-h withdrawal) and with dopaminergic medication (DOPA^ON^, after administration of a suprathreshold levodopa dose equal to the usual morning intake plus 50 mg). The first session occurred 1–4 days postoperatively, allowing coupling with MLR-LFP recordings. Two patients were unable to perform this postoperative session due to fatigue. The remaining four patients completed 153 trials, during which all exhibited freezing with 38 FOG episodes recorded (**Supplementary Table 1**). The following sessions were performed under CuN stimulation (DBS^CuN^), PPN stimulation (DBS^PPN^), or sham stimulation (DBS^OFF^), each applied for a 2-month period, in a cross-over randomized order with double-blind assessments (see ^13^).

Gait initiation was recorded using a 0.9 × 1.8 m force platform (AMTI, Watertown, MA, USA). Each trial began with quiet standing until a visual cue signaled the start of walking; patients then walked straight for 6–8 m, turned, and returned to the platform. Ground reaction forces and moments were sampled at 1000 Hz. Whole-body kinematics during straight walking and turning were acquired with a 32-marker motion capture system (Vicon®, Oxford, UK). Bilateral surface EMG recordings of the soleus and tibialis anterior muscles were sampled at 2000 Hz. The severity of axial motor symptoms was assessed using the MDS-UPDRS part III axial subscore (sum of items: speech, arising from chair, gait, freezing of gait, postural stability, posture) ^93^ and the Gait and Balance Scale (GABS) ^94^ (**Supplementary Table 1**).

### Gait initiation parameters and FOG events

Gait initiation was segmented into two phases: (1) anticipatory postural adjustments (APAs), from the initial biomechanical event (t□) to foot-off of the leading limb (FO□); and (2) step execution, from FO□ to the first foot contact (FC□). Seventeen spatiotemporal GI parameters were extracted for each trial from ground reaction forces and moments along the mediolateral (x), anteroposterior (y), and vertical (z) axes (**Supplementary Figure 1**). Trials in which t□ could not be reliably identified were excluded from analysis. FOG episodes were annotated from motion capture and EMGs recordings, defined as a transient inability to initiate or continue stepping which bursty co-contractions of antagonist muscles. Each episode was marked online during acquisition and subsequently verified offline by two independent raters. Gait initiation trials were classified as FOG+ (containing at least one FOG episode) or FOG-(without FOG).

### Gait initiation statistical analysis

The GI parameters extracted from force platform recordings were projected onto a varimax-rotated principal component analysis (RPCA) space obtained in a previous cohort of 38 PD patients performing a similar task ^48^. GI parameters were also compared to 24 age-matched healthy controls (HC) (see ^48^ for details) (**Supplementary Fig. 1**). In brief, we obtained 5 orthogonally RPC scores accounting for 73% of variance, each interpreted based on correlations with the original gait initiation parameters using Pearson’s correlation test. RPC1 related to forward stepping vigor and propulsion, which we labelled *Pace*; RPC2 reflected step times and cadence, which we labelled *Rhythm*; and RPC5 represented step width and lateral CoM speed, which we labelled *Lateral balance* (**Supplementary Fig. 1**)

We first examined how gait initiation performance scores changed during post-operative gait trials using linear mixed-effect models (LMM) with FOG imminence (FOG- vs. FOG+) and dopaminergic medication (DOPA^OFF^ vs. DOPA^ON^) as predictors with full interaction and patients as a random effect. Then, we explored how gait initiation RPC scores were modified across DBS conditions, using a LMM with dopaminergic medication (DOPA^OFF^ vs. DOPA^ON^) and DBS condition (DBS^OFF^ vs. DBS^CuN^ vs. DBS^PPN^) as predictors with full interaction and patients as random effect. We report estimated regression coefficient (β) ± standard error and associated *p*-values. All statistical analyses were performed in R (v4.2.2) using the *lme4* and *emmeans* packages for LMMs. Bonferroni correction was applied for multiple comparisons (N=5 RPCs), and corrected *p*-values (*p_c_*) < 0.05 were considered significant.

### MLR-LFPs recordings

MLR-LFPs were acquired using bipolar montages constructed from seven adjacent contact pairs of the DBS lead (Boston Scientific Cartesia; contact diameter 1.3 mm, height 1.5 mm, inter-contact spacing 0.5 mm; ventral contact labeled 1, dorsal contact labeled 8). LFPs were recorded continuously during gait initiation and walking, amplified, band-pass filtered (0.25–300 Hz), sampled at 512 Hz, and wirelessly transmitted via Bluetooth to a recording computer through a portable amplifier (Porti 32, TMS International, Enschede, The Netherlands) carried in a small backpack worn by the patient (Figure 1a). Kinetic signals from gait initiation and LFP activity were recorded simultaneously, synchronized, and exported for offline analysis using a MATLAB-based toolbox (MathWorks Inc.; http://biomechanical-toolkit.github.io). The implantable neurostimulator (Vercise PC, Boston Scientific) was placed the day after LFPs acquisition.

### LFPs data processing

MLR-LFPs were digitally high-pass filtered at 1 Hz, notch filtered at 50 Hz, and epoched from 2 s before the cue to the end of the second swing phase. Time–frequency decomposition was performed using a multitaper spectral estimation algorithm (Chronux toolbox, http://chronux.org/) with three orthogonal tapers and a 500 ms sliding window advanced in 30 ms steps. Power spectra (1–50 Hz) were baseline normalized to an 800 ms pre-trial period and expressed in decibels (10 × log□□). Time–frequency representations were screened for transient artifacts. Trials were excluded if mean power exceeded 5.7-fold baseline or if peak power exceeded 7.5 dB within –0.5 to +0.5 s from gait initiation onset (1–100 Hz range). In addition, all trials were visually inspected by at least two independent examiners for saturation, spectral smearing, and movement- or cable-induced artifacts.

### LFPs data statistical analysis

We examined changes in MLR-LFPs at gait initiation by applying a LMM at each point of the time-frequency map, using FOG imminence (FOG- vs. FOG+ trials), dopaminergic medication (DOPA^OFF^ vs. DOPA^ON^) and localization (PPN vs. CuN) as predictors with full interactions, and recording dipole nested within subjects as random effect. After identifying clusters of interest, we applied the same LMM in 500-ms time windows within defined frequency bands (alpha: 8-12 Hz; low-beta: 13-20 Hz; high-beta: 20-35 Hz). We then followed the same logic to examine, for each RPC, the relationship between MLR-LFPs and RPC scores. To do so, we added the RPC score as an interacting continuous predictor to the previous LMMs. We report the estimated regression coefficient (β) ± standard error and associated *p*-values. All statistical analyses were performed in R (v4.2.2) using the *lme4* and *emmeans* packages for LMMs. We corrected for multiple comparisons to control the false discovery rate and considered adjusted *p*-values < 0.05 as significant.

For regressions of post operative LFP power and *Rhythm* with DBS, we performed a linear regression with *Rhythm* score and localization/DBS condition (PPN vs. CuN) as fully interacting factors. A bonferroni correction was applied for multiple comparisons (N=5 RPCs), and corrected *p*-values (*p_c_*) < 0.05 were considered significant.

### Reporting summary

Further information on the research design is available in the Nature Portfolio Reporting Summary linked to this article.

### Data availability

Due to data protection regulations of Salpêtrière Hospital, INSERM and Paris Brain Institute, the raw gait and MLR LFPs data used in this study are available from the corresponding author (M.L.W) after approval of the IRB of these institutions. Source data are provided with this paper.

### Code availability

The custom codes used to generate the figures and statistics are available from the lead contact (M.L.W) upon request.

## Acknowledgments

The authors sincerely thank our patients for their dedication in participating in this research. This study was supported by the Institut National de la Recherche Médicale (INSERM) and the ‘Investissements d’avenir’ program (ANR-10-IAIHU-06 and ANR-11-INBS-0006) and grants from the Michael J Fox Foundation for Parkinson’s disease (grant number: 10019). We are also grateful to the nurses of the Clinical Investigation Center at the Paris Brain Institute for their assistance in organizing the research program, to Angele Van Hamme and Déborah Ziri for their help in data recordings.

A.C.C. receives funding from European Union’s Horizon 2020 research and innovation program under the Marie Skłodowska-Curie grant (H2020-MSCA-IF-2019, grant agreement No 898265) and the Swiss National Science Foundation (SNSF); M.Y. receives support by the Paris Sorbonne University; Y.M receives funding from the European Union’s Horizon 2020 research and innovation program under the Marie Slodowska-Curie-Horizon 2020 (H2020-MSCA-IF-2019, grant agreement No 898265). B.L. receives funding from the Agence Nationale de la Recherche (Grant No. R16150DD); M.L.W receives funding from the Agence Nationale de la Recherche (CoEN program), Fondation de France, France Parkinson, and Boston Scientific.

## Authors contributions

M.L.W, C.K, B.L. designed the research; M.Y, A.C.C, J.B., D.G., C.O, H.B, C.K., and M.L.W. acquired the data; M.Y, A.C.C, K.L., J.B., Y.M., N.G. ,D.G., H.T., M.R., C.O., H.B., B.L., C.K. and M.L.W. made the analysis and interpreted the data; M.Y, A.C.C, C.K., B.L. and M.L.W. made the first draft of manuscript; M.Y, A.C.C, C.K., B.L. and M.L.W. performed revision of the manuscript

