## Supplemental Table 1, Supplemental Figures 1-5 for "Distinct roles of the human cuneiform and pedunculopontine nuclei in gait initiation and freezing of gait"

**Supplementary Materials**

| **Patient** | **P01** | **P02** | **P03** | **P04** |
| --- | --- | --- | --- | --- |
| Age, y/Sex | 65/M | 68/M | 67/F | 54/M |
| Disease duration, y | 6 | 12 | 10 | 12 |
| MDS-UPDRS part 3, OFF/ON | 29/17 | 37/19 | 51/28 | 27/12 |
| Axial score, OFF/ON | 7/6 | 9/7 | 16/11 | 6/3 |
| GABS, OFF/ON | 10/12 | 29/24 | 40/25 | 11/4 |
| FOG-Q | 29 | 36 | 36 | 33 |
| MDRS | 144 | 142 | 141 | 139 |
| LEDD (mg/d) | 650 | 2,200 | 860 | 1,456 |

**Supplementary Table 1. Demographic and clinical characteristics of 4 PD patients operated for MLR DBS**

FOG-Q: Freezing of Gait-questionnaire; GABS: Gait and balance scale; LEDD: levodopa equivalent daily dosage; MDS-UPDRS: Movement Disorders Society-Unified Parkinson’s Disease Rating Scale; OFF/ON: without (OFF) and with (ON) dopaminergic treatment.

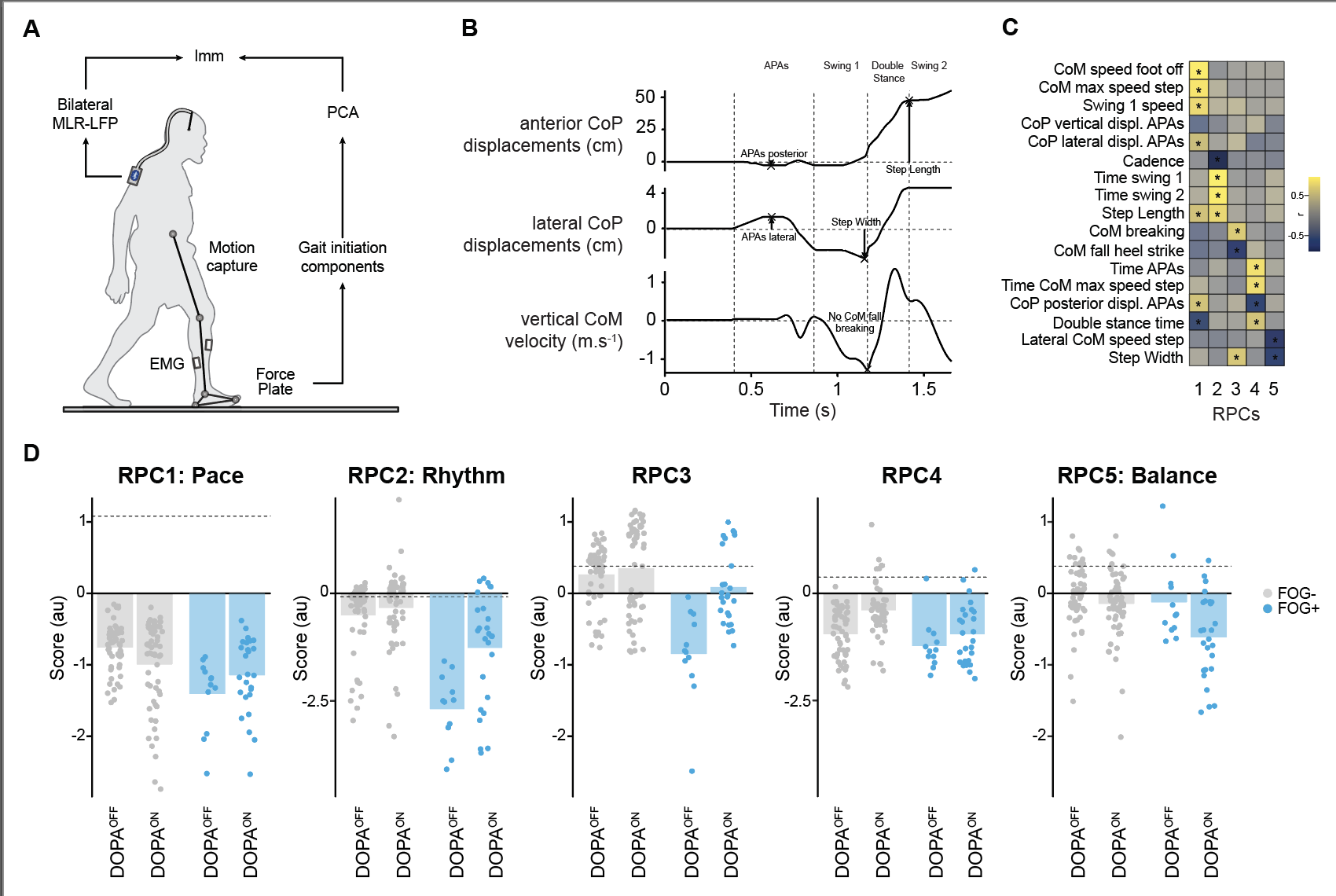

**Supplementary Fig. 1 | Gait initiation kinematic parameters**

**a**. MLR LFP recordings synchronized with gait initiation recordings

**b.** Gait initiation kinematic parameters extracted from the force platform. APAs: anticipatory postural adjustments. CoP: center of foot pressure, CoM: center of mass

**c**. Rotated principal component analysis including the 17 gait initiation parameters. displ: displacement

**d**. RPC scores without (DOPAOFF) and with (DOPAON) dopaminergic medication for FOG- (grey) and FOG+ (blue) trials. Dashed lines: mean scores obtained in healthy controls, zero lines: mean scores obtained in a previous cohort of 38 PD patients with dopasensitive FOG.

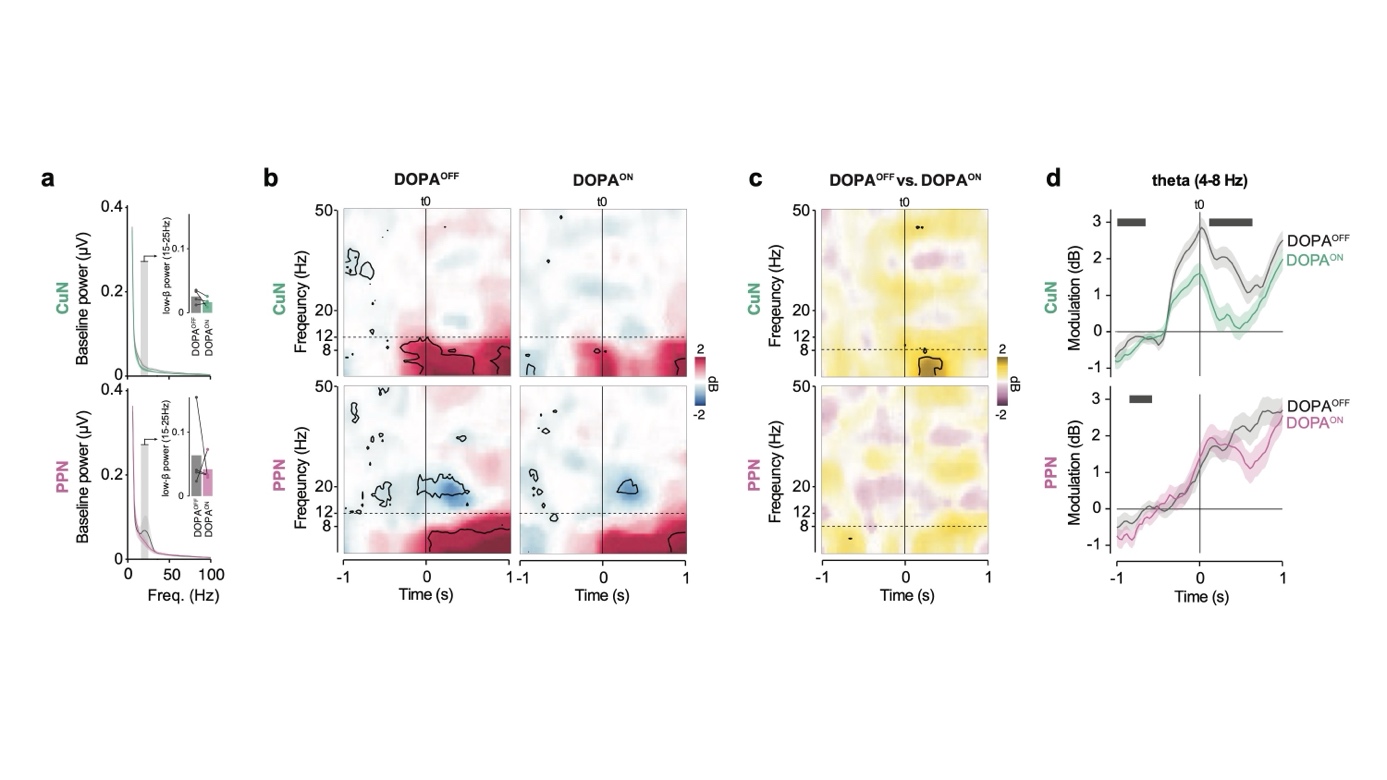

**Supplementary Fig. 2 | Effects of dopaminergic treatment on CuN and PPN neural activity at gait initiation**

**a**. Mean power spectral density across patients at baseline in the CuN and PPN, and DOPA^OFF^ and DOPA^ON^. Focus on average baseline power in the beta (15–25 Hz) band. Dots represent patient-level averages across contacts. No statistical effect observed.

**b**. Time–frequency maps showing LFPs modulations relative to baseline in the CuN and PPN, and DOPA^OFF^ and DOPA^ON^. Dashed line: APAs onset. Black contours: p < 0.05, LMM.

**c**. Time–frequency differences in LFPs modulation between DOPA^OFF^ and DOPA^ON^ separately for PPN and CuN. Black contours: p < 0.05, LMM.

d. Temporal profiles of theta (4-8 Hz) power modulation DOPA^OFF^ and DOPA^ON^ in CuN (green) and PPN (purple). Black bars indicate significant differences between DOPA^OFF^ and DOPA^ON^ conditions (p < 0.05, LMM). Shaded areas represent SEM.

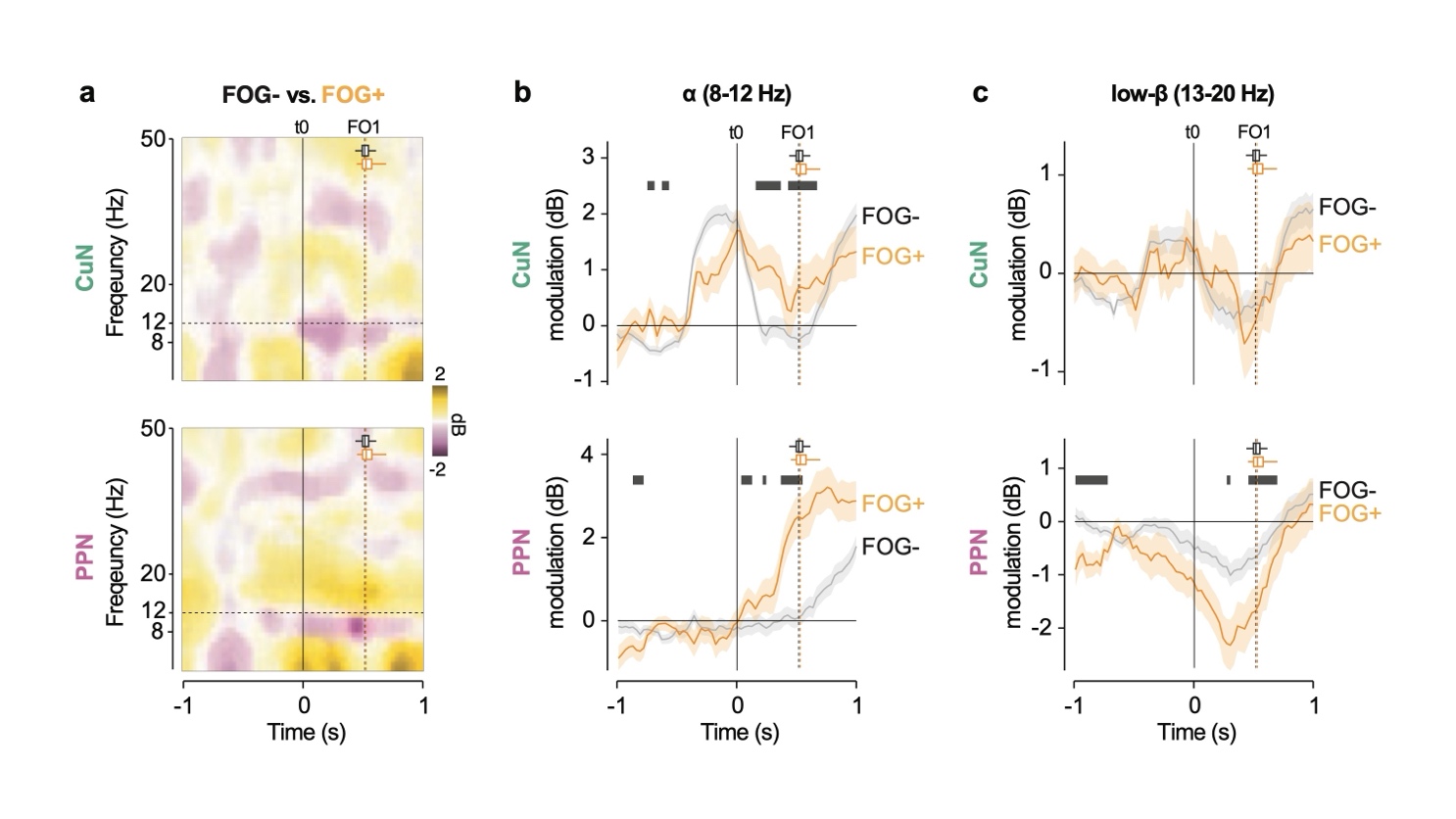

**Supplementary Fig. 3 | Effects of FOG imminence on MLR LFP modulation at gait initiation**

**a.** Time–frequency maps showing LFPs modulations relative to baseline for FOG- and FOG+ trials. Black contours: p < 0.05, LMM.

**b**. Time–frequency differences in LFPs modulation between FOG- and FOG+ trials. Black contours: p < 0.05, LMM.

**c**. Alpha and beta LFPs power modulation in for FOG- (grey lines) and FOG+ (orange lines) trials. Black bars: p < 0.05, LMM.

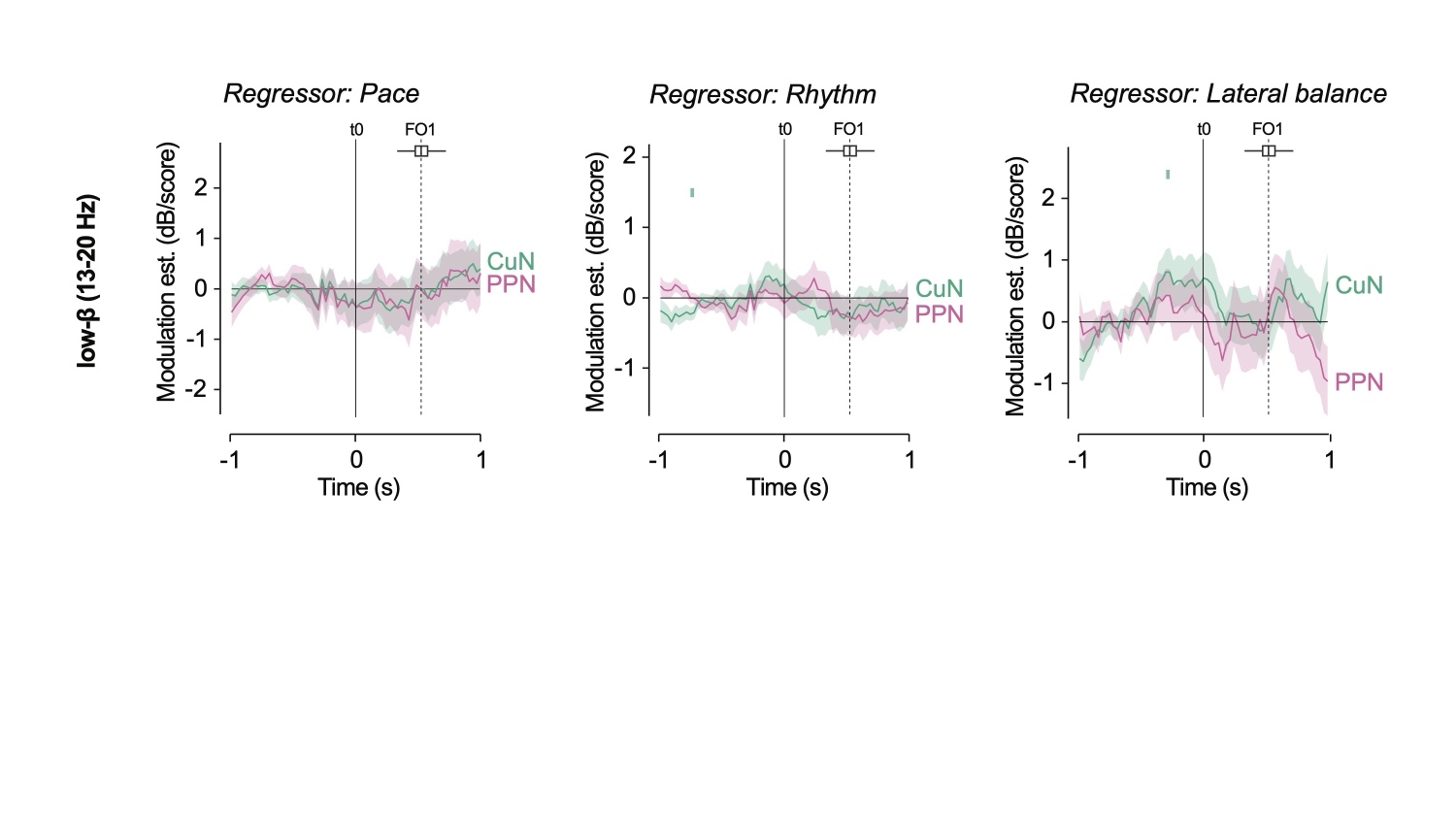
 **Supplementary Fig. 4 | Correlation between low-beta MLR LFP modulation and gait initiation components**

Graphs report the relationship between low-beta band CuN (green curves) and PPN (purple curves) LFP modulation at gait initiation and *Pace*, *Rhythm*, and *Lateral balance* scores. Black bars: p < 0.05 between CuN and PPN, green and purple bars: p < 0.05 for correlation analysis in CuN and PPN, respectively, LMM. FO1= foot-off, t0= first biomechanical event, i.e. anticipatory postural adjustment phase start.

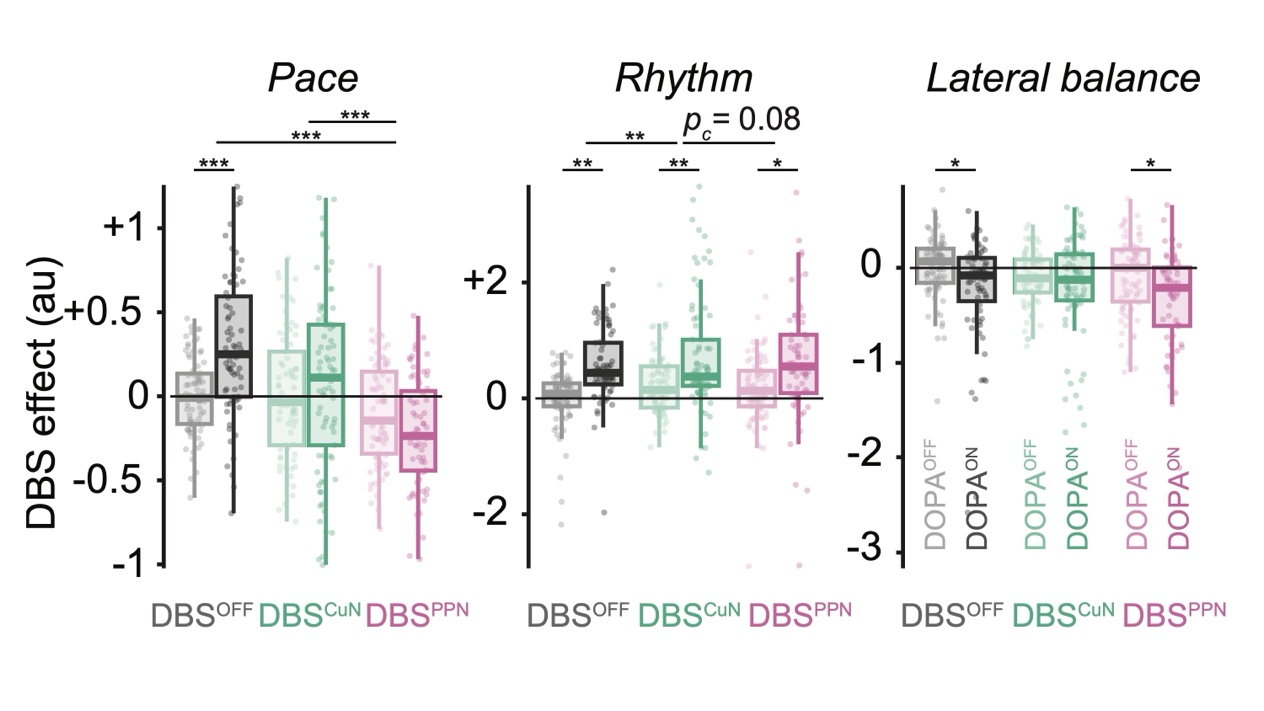
**Supplementary Fig. 5 | Effects of MLR-DBS on gait initiation without and with dopaminergic medication**

Changes in RPC scores, *Pace*, *Rhythm* and *Lateral balance*, without (OFF_Sham, grey) and with CuN (green) or PPN (purple) DBS, without (DOPA^OFF^, light) and with dopaminergic medication (DOPA^ON^). *p < 0.05, **p < 0.01, ***p<0.001, LMM.
